# A new demethylase gene *OsDML4* involved in high temperature induced floury endosperm formation in rice (*Oryza sativa* L.)

**DOI:** 10.1101/2022.01.20.477110

**Authors:** Yan Yan, Chao Li, Zhen Liu, Jun-Jie Zhuang, Jia-Rui Kong, Zhen-Kun Yang, Jie Yu, Mohammad Shah Alam, Cheng-Cheng Ruan, Heng-Mu Zhang, Jian-Hong Xu

## Abstract

High temperature (HT) can affect the accumulation of seed storage materials and cause adverse effects on the yield and quality in rice. DNA methylation plays an important role in plant growth and development. However, the temperature and DNA methylation interaction on rice seed development has not been studied yet. Here, we identified a new demethylase gene *OsDML4* and discovered its function on cytosine demethylation to affect the endosperm formation during the grain filling. Knockout of *OsDML4* induced floury endosperm only under HT, which resulted from dramatically reduced the transcription and accumulation of glutelins and 16-kDa prolamin. The expression of two important transcription factors *RISBZ1* and *RPBF* was significantly declined in the *osdml4* mutants. The absence of *OsDML4* also caused adverse effects on the formation of protein bodies (PBs), the number of PB-II was greatly decreased and incomplete PB-II with empty space and abnormally shaped PB-II were observed in the *osdml4* mutants. Whole-genome bisulfite sequencing analysis of seeds at 15 days after pollination revealed much higher global methylation levels of CG, CHG and CHH contexts in the *osdml4* mutants compared to wild type (WT). Moreover, the methylation status of *RISBZ1* promoter was hypermethylated but *RPBF* promoter was nearly unchanged. No significant difference was detected between WT and the *osdml4* mutants under room temperature. In conclusion, our study demonstrates a novel *OsDML4*-mediated epigenetic regulatory mechanism involving in the formation of floury endosperm, which will provide a new perspective in regulating endosperm development and the accumulation of SSPs in rice.

## Introduction

Rice (*Oryza sativa* L.) is one of the most important staple crops in the world, feeding more than 60% of the population in China and providing around 40% of the total calorific needs (Cheng et al., 2007). With the continuous global warming and the emergence of extreme weather more frequently, the yield and quality of rice are more and more seriously threatened (Peng et al., 2004; Lin et al., 2005). The growth and development of rice during the grain filling stage is greatly affected by temperature. High temperature (HT) can increase grain chalkiness and total protein content, reduce grain weight, total starch content and amylose content, and affect the expression of a series genes related to the biosynthesis of seed storage materials thus leading to adverse influences on the yield and quality of rice (Yamakawa et al., 2007; Lin et al., 2010; Li et al., 2011; Cao et al., 2017; Tabassum et al., 2020; Xu et al., 2020).

Seed storage proteins (SSPs) are the important component in rice grain, which provide the nitrogen source for seed germination and nutrients for human and livestock (Kawakatsu and Takaiwa, 2010). SSPs can be divided into glutelins, prolamins, globulins and albumins according to the different solubility (Shewry and Casey, 1999). Prolamins are the major component of SSPs in most cereals such as maize (*Zea mays*), wheat (*Triticum aestivum*) and barley (*Hordeum vulgare*) (Shewry and Tatham, 1999). While glutelins account for the highest proportion of SSPs in rice (Takaiwa, 1999). Based on molecular mass, glutelins can fall into 57-kDa proglutelin, 37-kDa acid subunit and 22-kDa basic subunit (Yamagata et al., 1982), and fifteen glutelin genes have been identified in the rice reference genome, which can be classified into four subfamilies (*GluA*, *GluB*, *GluC* and *GluD*) (Kawakatsu et al., 2008). The prolamins fall into three groups: 10-, 13- and 16-kDa (Ogawa et al., 1987), and a total of 34 prolamin genes are present in the rice reference genome (Xu and Messing, 2009). The temperature has a great effect on the accumulation of SSPs and the expression of SSP biosynthesis genes. The HT could reduce the content of 13-kDa prolamin and the expression of 13-kDa prolamin genes, while the expression of glutelin genes was less affected (Yamakawa et al., 2007). Whereas, the HT can also increase the glutelin content and decrease the prolamin content (Ashida et al., 2013; Cao et al., 2017).

SSPs are synthesized on the rough endoplasmic reticulum (rER) and form two types of protein bodies (PBs) in rice endosperm cells. Spherical protein body I (PB-I) is formed by prolamins, which are retained in the ER lumen after synthesis, whereas irregular shaped protein body II (PB-II) is formed by glutelins that are transported to the protein storage vacuoles (PSVs) via the Golgi apparatus (Tian and Okita, 2014; Tian et al., 2018). Furthermore, many important regulatory factors were involved in protein folding process and the formation of PBs. *OsVPE1* is essential for glutelin maturation, and the mutation of *OsVPE1* influences the shape of PB-II from round to irregular (Wang et al., 2009). An ER luminal binding protein (*BIP*) is involved in folding and assembly of prolamin. Overexpression of *BiP1* resulted in floury and shrunken endosperm with a significant reduction of SSPs and altered the morphology of PB-I (Yasuda et al., 2009). Protein disulfide isomerase (PDI) is required in glutelin trafficking and plays a critical role in proglutelin maturation and segregation of proglutelin and prolamin with the ER lumen. The absence of PDI induces floury endosperm and small ER-derived PBs containing both proglutelin precursor and prolamin (Takemoto et al., 2002; Han et al., 2012).

The endosperm specific transcription factors (TFs) have been demonstrated to regulate the expression of SSP biosynthesis genes. The basic leucine zipper factor *RISBZ1*/*OsbZIP58* and rice prolamin binding box (*RPBF*) are two important regulators in rice, which can bind to specific motifs including GCN4, prolamin box (P box), ACGT and AACA in the promoter regions of SSP biosynthesis genes and activate their expression (Takaiwa et al., 1996; Wu et al., 2000; Yamamoto et al., 2006; Kawakatsu et al., 2009). Moreover, *RISBZ1* can inhibit the expression of SSP genes by inducing its alternative splicing (Xu et al., 2020). In addition, *OSMYB5* functions as trans-acting regulator for glutelin genes through binding to the AACA motif (Suzuki et al., 1998). Little is known about the expression patterns of SSP biosynthesis genes and the mechanism of corresponding regulatory factors under HT in rice.

DNA methylation plays a critical role in many biological processes such as embryonic development, gene regulation, structural stability of chromatin, and various biotic and abiotic stress responses (Bird, 2002; Bender, 2004; Zhang et al., 2018). In plants, DNA methylation occurs at cytosine in CG, CHG and CHH sequences, and methylated DNA can be removed by demethylation (Penterman et al., 2007; Zhu, 2009). Six demethylase genes have been identified in rice including four *REPRESSOR OF SILENCING1*(*ROS1*) ortholog genes and two *DEMETER-LIKE3* ortholog genes (*DML3*) (Zemach et al., 2010). Knock-in null mutation of *OsROS1a* leads to abortion of early-stage endosperm development, formation of irregular embryos and production of no seeds (Ono et al., 2012). Knockout or knockdown of *DNG701*(*OsROS1b*) can increase DNA methylation level and thus inhibit the expression of the retrotransposon *Tos17* (La et al., 2011). *OsROS1a*-mediated DNA demethylation can change the number of aleurone layers and improve nutritional value of rice grains (Liu et al., 2018). Hypomethylation by *OsROS1a* in rice vegetative cells increases DNA methylation in sperm (Kim et al., 2019). *DML3*-Mediated DNA demethylation can delay leaf senescence in *Arabidopsis* (Yuan et al., 2020).

A new demethylase gene named *OsDML4* was identified based on conservative amino acid sequences of glycosylase domain, which is highly expressed during the reproductive period (Liu et al., 2014). In order to understand whether *OsDML4* may have function in DNA demethylation, the knockout mutants of *OsDML4* were obtained using CRISPR-Cas9 genome editing system, which showed the phenotype of increased grain chalkiness and floury endosperm under HT and greatly increased the methylation levels in CG, CHG and CHH of the whole genome only under HT. These results showed that the new demethylase gene *OsDML4* was involved in the formation of floury endosperm through epigenetic regulatory mechanism depended on temperature in rice.

## Results

### Grain appearance of the *osdml4* mutants

In order to investigate the function of *OsDML4*, the CRISPR/Cas9 system was used to generate two frameshift *OsDML4* knockout mutants (Figure 1). Under HT conditions, the seeds of *osdml4* mutants appeared chalky and floury in cross-section, while the WT seeds are transparent (Figure 2A-B). Scanning electron microscopy (SEM) analysis revealed that the starch granules of the *osdml4* mutants were round in shape and loosely packaged, whereas those of WT are polygons and tightly packaged (Figure 2C). These morphological changes in the starch granules may account for the floury features of the *osdml4* mutants to some extent, indicating that the grain filling process might be damaged in the *osdml4* mutants. The storage materials in the seeds were then examined to find that the total protein content was increased and the total starch content was decreased in the *osdml4* mutants (Figure 2D-E), and the amylose content of the *osdml4* mutants exhibited a significant reduction compared to WT (Figure 2F). In addition, the seed length of the *osdml4* mutants was significantly longer than those of WT, while the thicknesses and 1000-grain weight were considerably reduced (Figure 3A-B, D-E). No obvious difference of seed width was observed between WT and the *osdml4* mutants (Figure 3C), and the main agronomic traits, with respect to plant height, effective tiller number, seed setting rate and seed number per panicle were not significantly changed (Figure S1).

**Figure 1.**
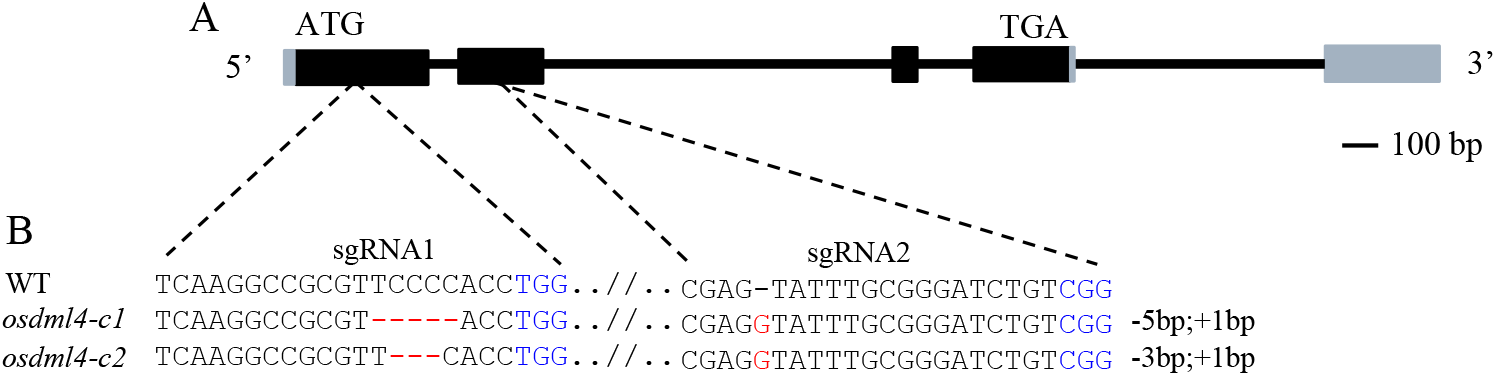
CRISPR/Cas9-induced mutations in the *OsDML4* gene. (A) The Schematic diagram of the *OsDML4* gene. The UTRs, exons and introns are indicated by gray rectangles, black rectangles and black lines, and the start codon (ATG) and stop codon (TGA) and their positions were showed. (B) The sequences of the two targets are shown with the protospacer adjacent motif (PAM) sequences labeled in blue color. The editing genotypes are identified by sanger sequencing and aligned with wild type (WT), the deletions and insertions are indicated by red dashes red letters.

**Figure 2.**
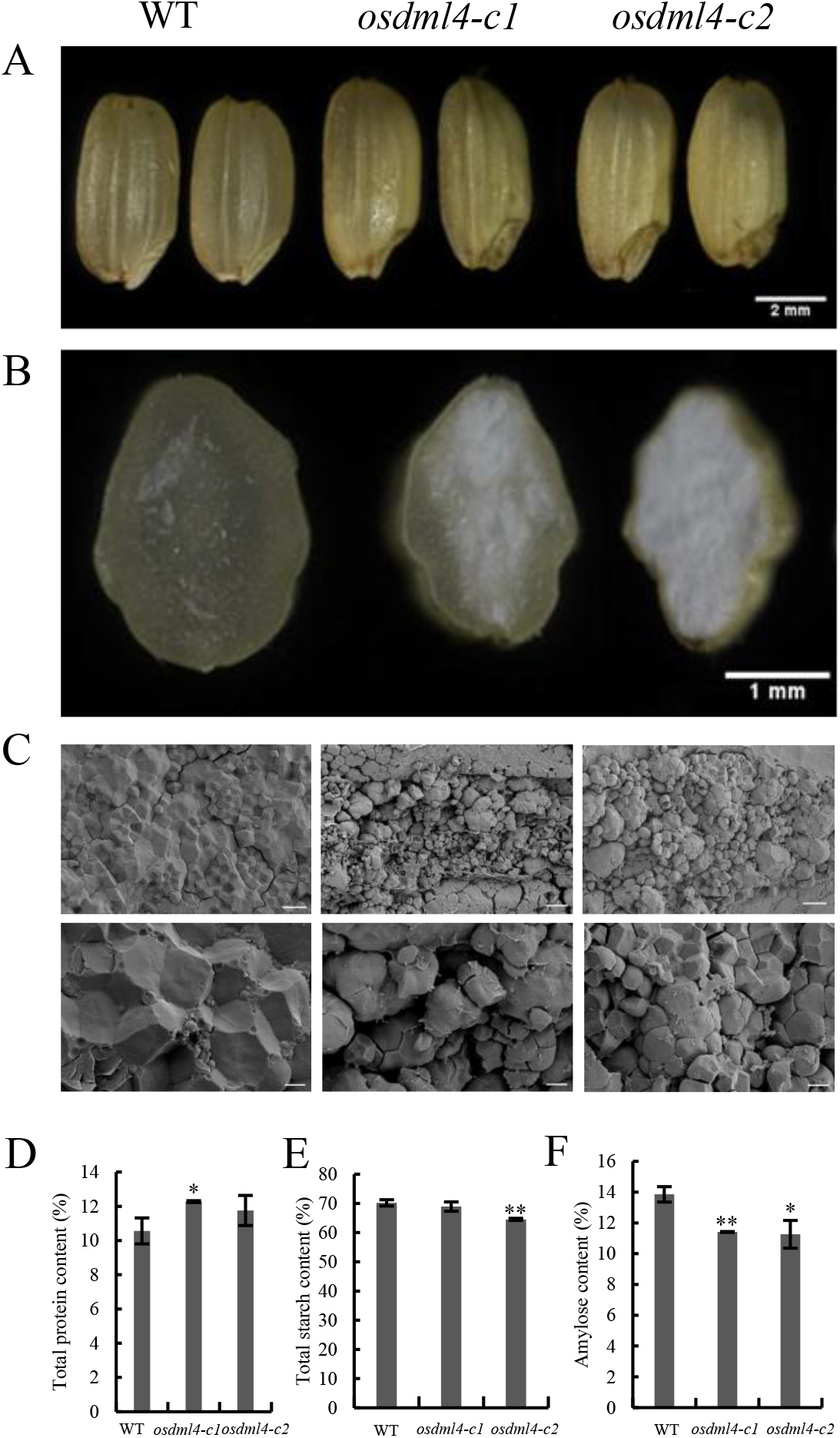
Appearance of the WT and the *osdml4* mutants mature seeds under HT. (A) External appearance of brown seeds from WT and the *osdml4* mutants. (B) Transverse sections of WT and the *osdml4* mutant dry seeds. (C) Scanning electron micrographs of mature endosperms of WT and the *osdml4* mutants (up, scale bar = 10μm ; down, scale bar = 3μm). (D) Total protein content in mature seeds of WT and the *osdml4* mutants. (E) Total starch content in mature seeds of WT and the *osdml4* mutants. (F) Amylose content of WT and the *osdml4* mutants. Data are the means ± SD of three biological replicates. Significant differences were determined using two-tailed Student’s t-test (*p<0.05, **p<0.01).

**Figure 3.**
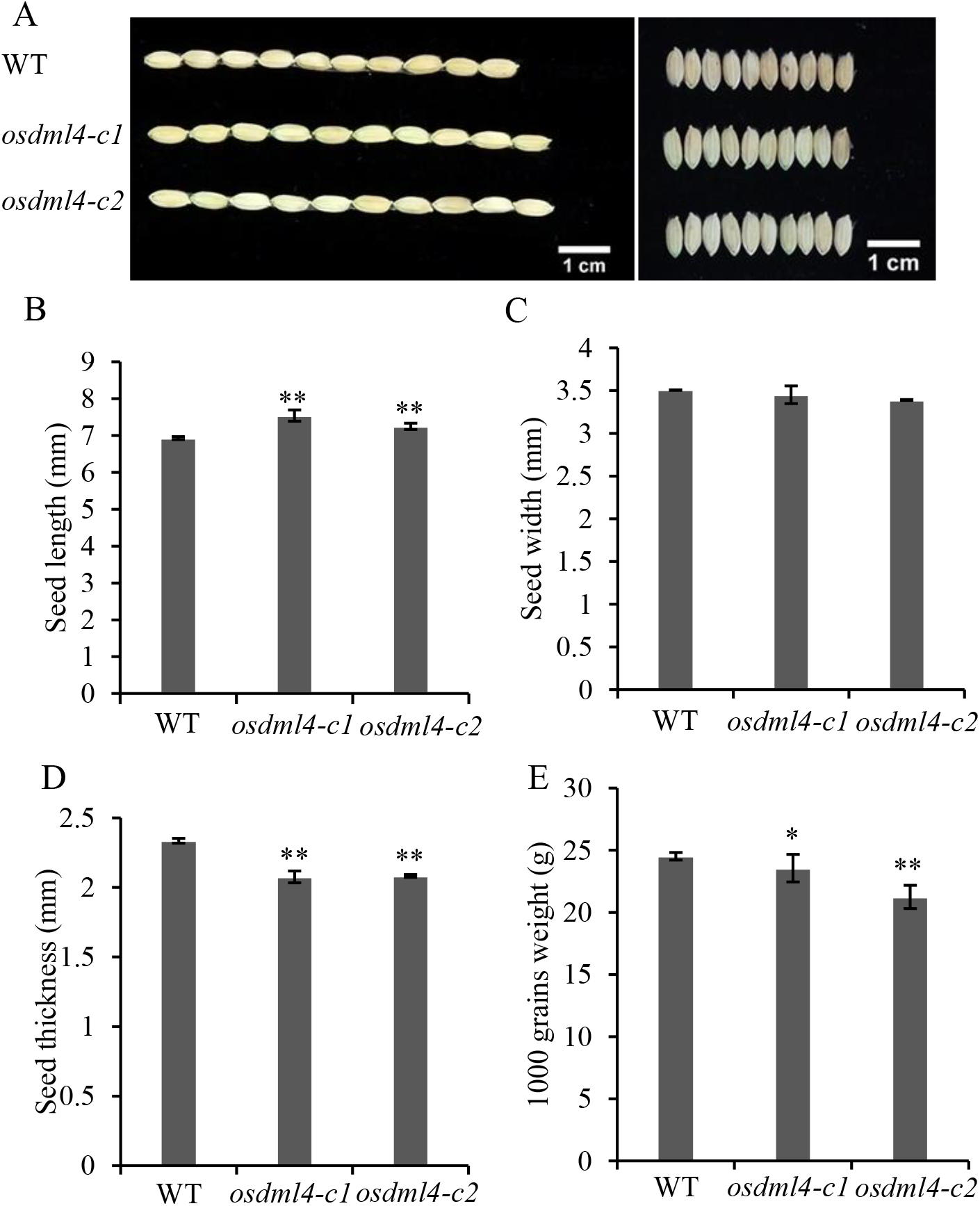
Grain shape analyses of the *osdml4* mutants under HT. (A) The Seed phenotype of the wild type and *osdml4* mutants. The statistic analyses of (B) seed length, (C) seed width, (D) seed thicknesses, and (E) 1000-grain weight of WT and *osdml4* mutants. Data are the means ± SD of three biological replicates. Significant differences were determined using two-tailed Student’s t-test (*p<0.05, **p<0.01).

As the temperature can affect the grain phenotype during the filling stage, the WT and the *osdml4* mutants were then grown under RT during the filling period. The results showed that the *osdml4* mutants showed similar phenotype to WT almost without chalky appearance of grain and the floury endosperm under RT (Figure S2). These results indicated that knockout of *OsDML4* can result in a floury endosperm and affect the normal close packaging of starch and modulate seed size only under HT.

### Knockout of *OsDML4* affects the accumulation of SSPs under HT

SDS-PAGE analysis revealed that the *osdml4* mutants contain much fewer amounts of 57-kDa proglutelin than WT under HT, accompanied by a remarkable decrease in the 40-kDa acidic and 20-kDa basic subunits of the mature glutelins (Figure 4A). In addition, the 16-kDa prolamin and 10-kDa prolamin was a little reduced, while the 13-kDa prolamin were considerably unchanged in comparison with WT (Figure 4B). While there had no significant difference in the contents of glutelin and prolamin in the mature seeds between the *osdml4* mutant and WT under RT (Figure 4C-D), suggested that knockout of *OsDML4* influences the normal storage protein inclusions to form a chalky endosperm during the maturation process of the seeds only under HT.

**Figure 4.**
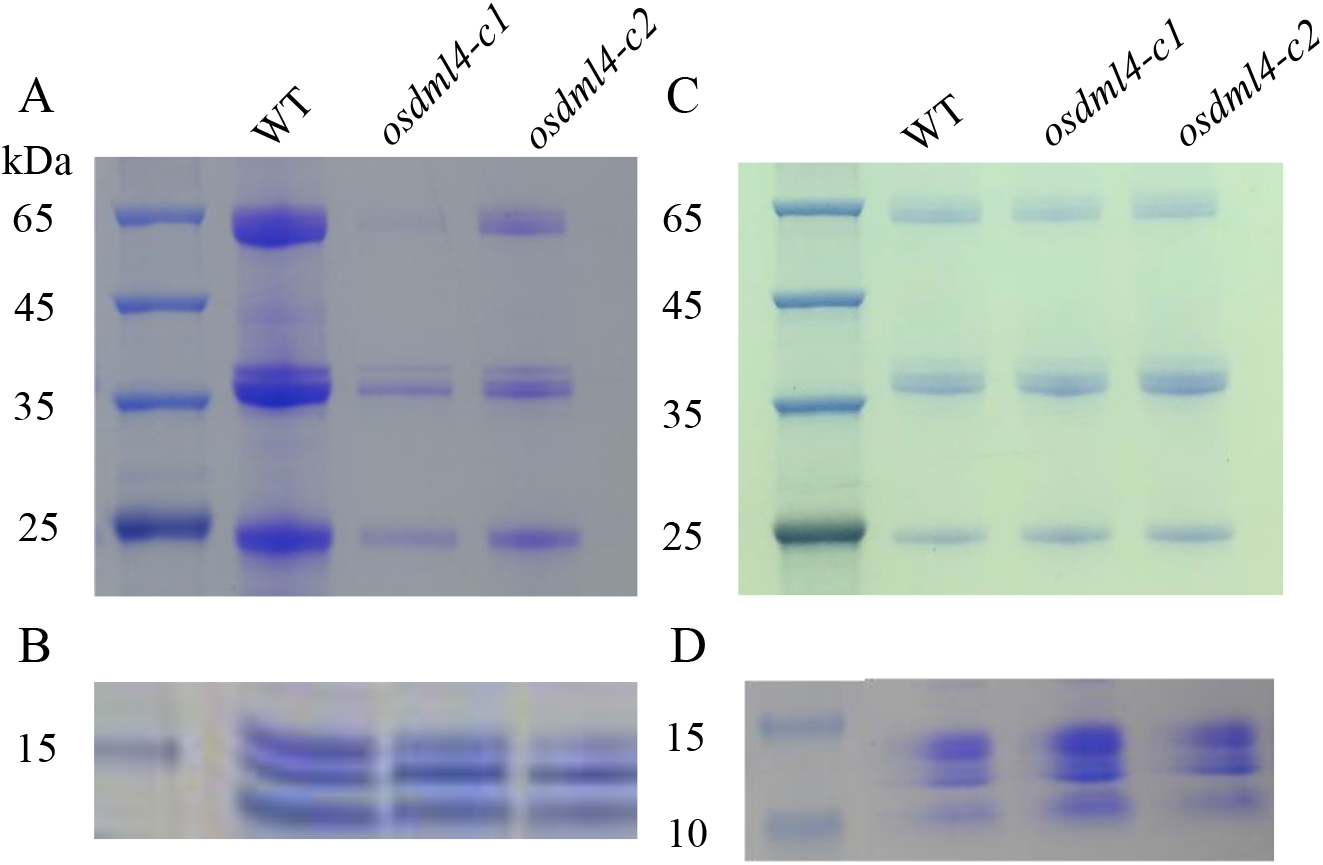
SDS-PAGE analysis of seed storage proteins in mature seeds of WT and the *osdml4* mutants. (A) glutelins and (B) prolamins in WT and *osdml4* mutants under HT. (C) glutelins and (D) prolamins in WT and *osdml4* mutants under RT. The seed storage proteins were extracted according to the previous method, and were separated by 12% SDS-PAGE.

### *OsDML4* affects the expression of SSPs biosynthesis genes and related TFs under HT

To determine whether the alterations of the SSPs accumulation in *osdml4* mutants will also be reflected at the transcriptional level, qRT-PCR was used to determine the expression levels of SSP biosynthesis genes. Under HT conditions, the expression of all glutelin genes were significantly suppressed in the *osdml4* mutant. The 10-kDa and 16-kDa prolamin genes were also greatly down regulated and the 13-kDa prolamin genes showed little difference between the *osdml4* mutants and WT (Figure 5A). Previous studies have shown that *RPBF* and *RISBZ1* are two important TFs that can positively activate the expressions of prolamin and glutelin genes. Therefore, the expression levels of both TFs were investigated, and found that their expression levels were significantly declined in the *osdml4* mutants compared to WT under HT, but no significant difference under RT (Figure 5 B and S3B). The transcription levels of genes related to SSPs proper folding and assembly in the post-translation process were also examined. The expression of *VPE1*, *PDIL1-1* was significantly repressed in the *osdml4* mutant under HT (Figure 5B). Under RT conditions, the glutelins and prolamins exhibited similar expression levels between the *osdml4* mutant and WT (Figure S3A). These results indicated that *OsDML4* could play a crucial role in the expression of SSPs biosynthesis genes under HT.

**Figure 5.**
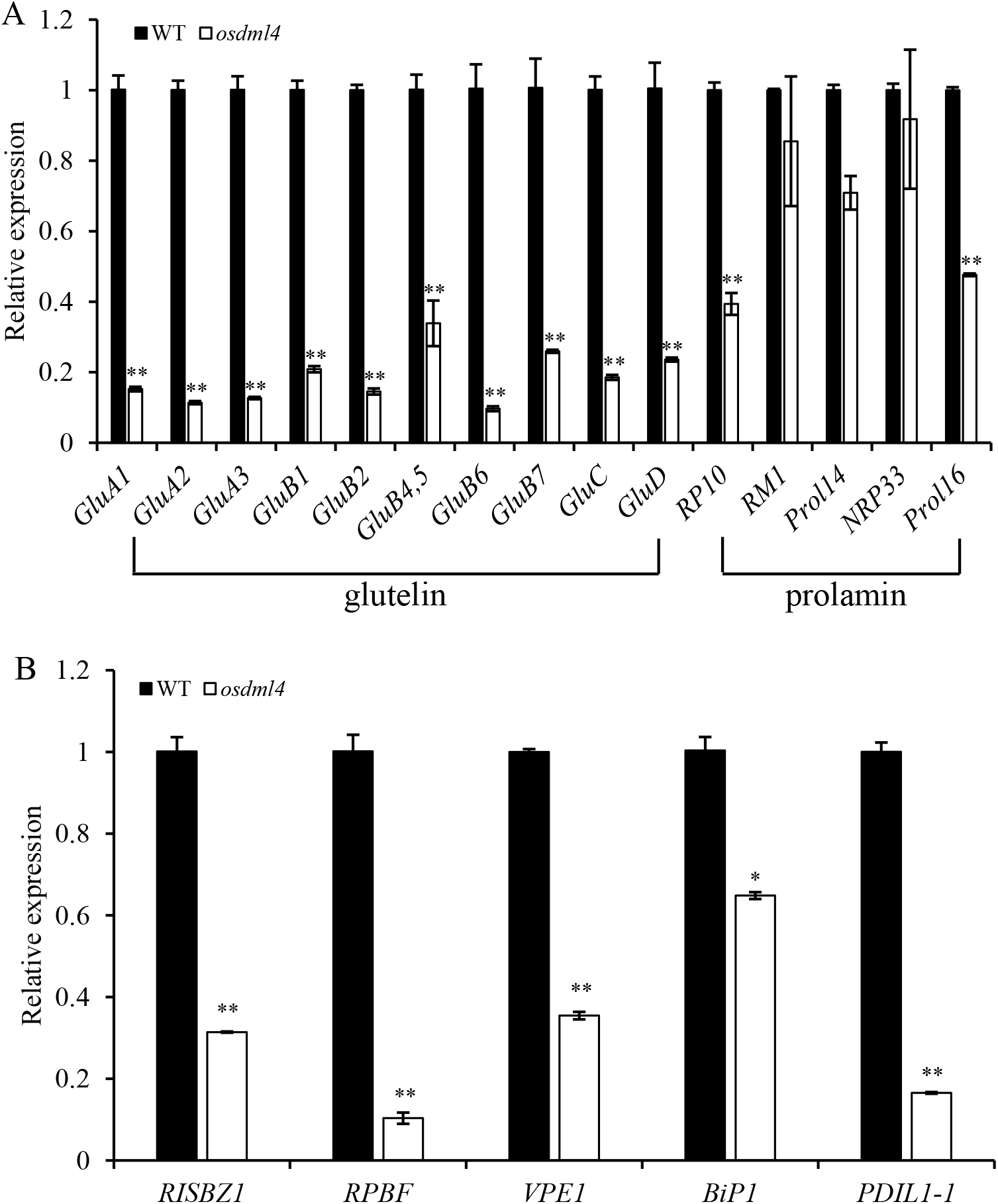
qRT-PCR analysis of SSP genes and their regulatory factors in 15 DAP immature seeds of WT and the *osdml4* mutant under HT. (A) SSP genes, (B) SSP regulatory factors. Data are the means ± SD of three biological replicates. Significant differences were determined using two-tailed Student’s *t*-test (*p<0.05, **p<0.01).

Because the profile of SSP was greatly altered in the *osdml4* mutants, the intracellular structures of developing endosperms at 15 DAP was observed by TEM to determine whether this mutation also influences the PB formation. Under HT, the two types of PBs were readily discernible in WT. Prolamin-containing PB-I were round and surrounded by ER, while glutelin-containing PB-II were larger and irregularly shaped (Figure 6A-C). In the *osdml4* mutant, the number of total PBs especially PB-II was greatly reduced (Figure 6D). In addition to the normal PB-II, some incomplete PB-II with empty space and small abnormally shaped PB-II were also observed in the *osdml4* mutant (Figure 6E, F). Furthermore, much smaller PB-Is were also observed and some of which were attached to abundant vesicular structures (Figure 6F). Whereas the size, shape and number of both PB-I and PB-II were similar to those of the WT under RT (Figure S4). These results illustrated that loss function of *OsDML4* has a great impact on PB formation and probably leads to the formation of floury endosperm under HT.

**Figure 6.**
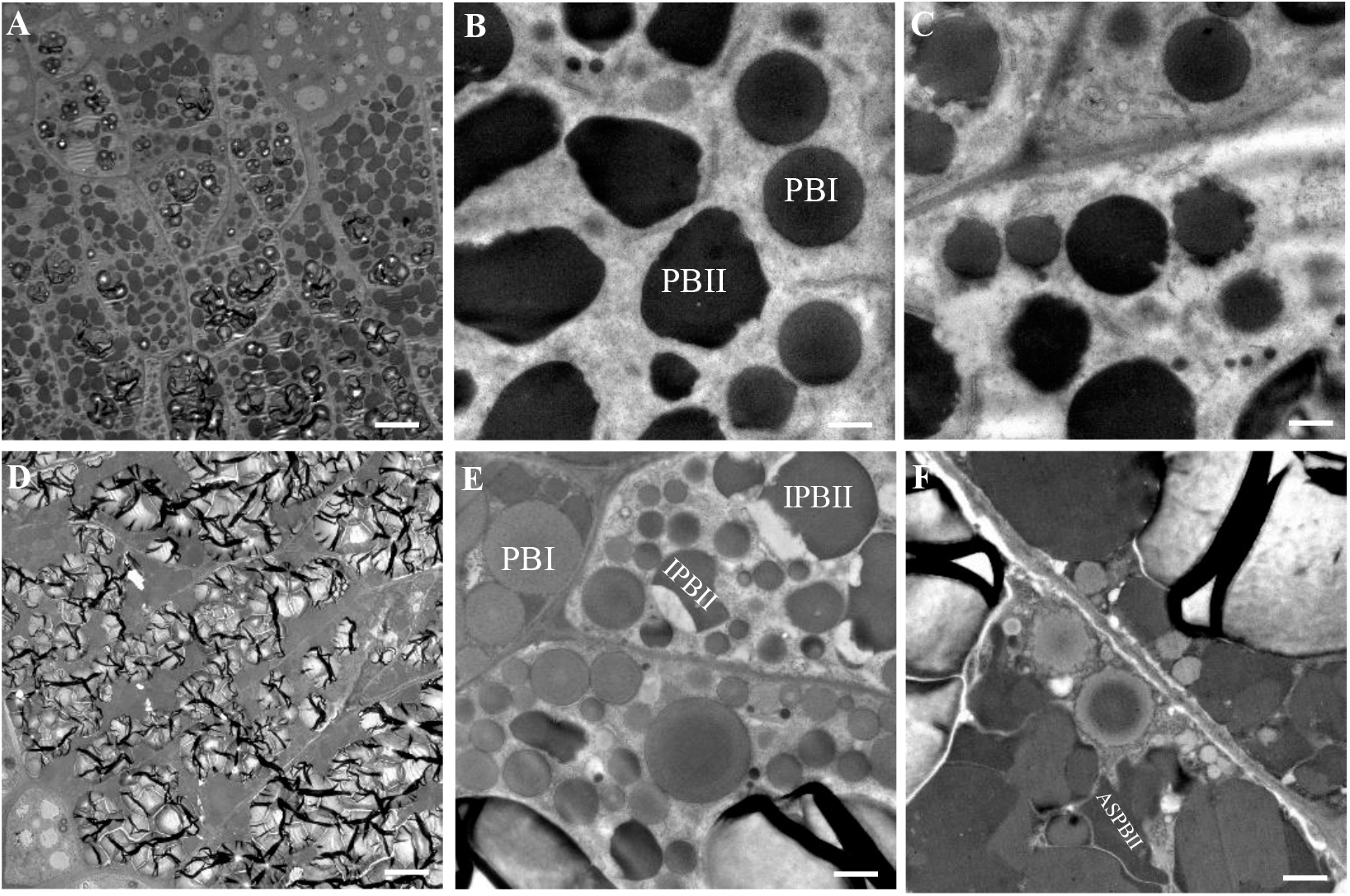
Transmission electron microscope of protein bodies in developing endosperm at 15 DAP under HT. (A-C) WT, (D-F) the *osdml4* mutant. Scale bar = 10 μm in (A, D) and 1μm in (B, C, E, F). IPBII, incomplete PBII with empty space; ASPBII: abnormally shaped PBII.

### Genome-wide hypermethylation in developing seeds of *osdml4*

Since *OsDML4* is a new identified demethylase gene in rice, to investigate whether *OsDML4* has the function of DNA demethylation, WGBS was carried out to set up the methylomes for 15 DAP seeds of WT and the *osdml4* mutants. The methylome covers more than 98% of all the genomic cytosine positions with >20-fold coverage per strand (Table S2). Comparative analysis of methylation levels between WT and the *osdml4* mutants showed much higher global methylation levels of CG, CHG and CHH contexts in the *osdml4* mutants than in WT under HT. While under RT, CG methylation levels in the *osdml4* mutants were slightly lower than that of WT, and CHG and CHH methylation levels were slightly increased in the *osdml4* mutants compared to WT (Figure 7A). We then analyzed the distribution of DNA methylation levels in different regions including promoter, 5’-UTR, exon, intron and 3’-UTR, and found that the global methylation levels of all three contexts (CG, CHG and CHH) in the *osdml4* mutants were sharply increased under HT. Interestingly, under RT, the *osdml4* mutants had moderately lower CG methylation levels compared with WT, while CHG methylation levels were almost identical to that of WT. On the contrary, the CHH methylation level was increased at the promoters, but the degree of increase was much less than that under HT (Figure 7B-C). These results suggested that *OsDML4* has the function of demethylation in all three contexts under HT, and loss function of *OsDML4* may result in the down regulation of genes associated with DMRs.

**Figure 7.**
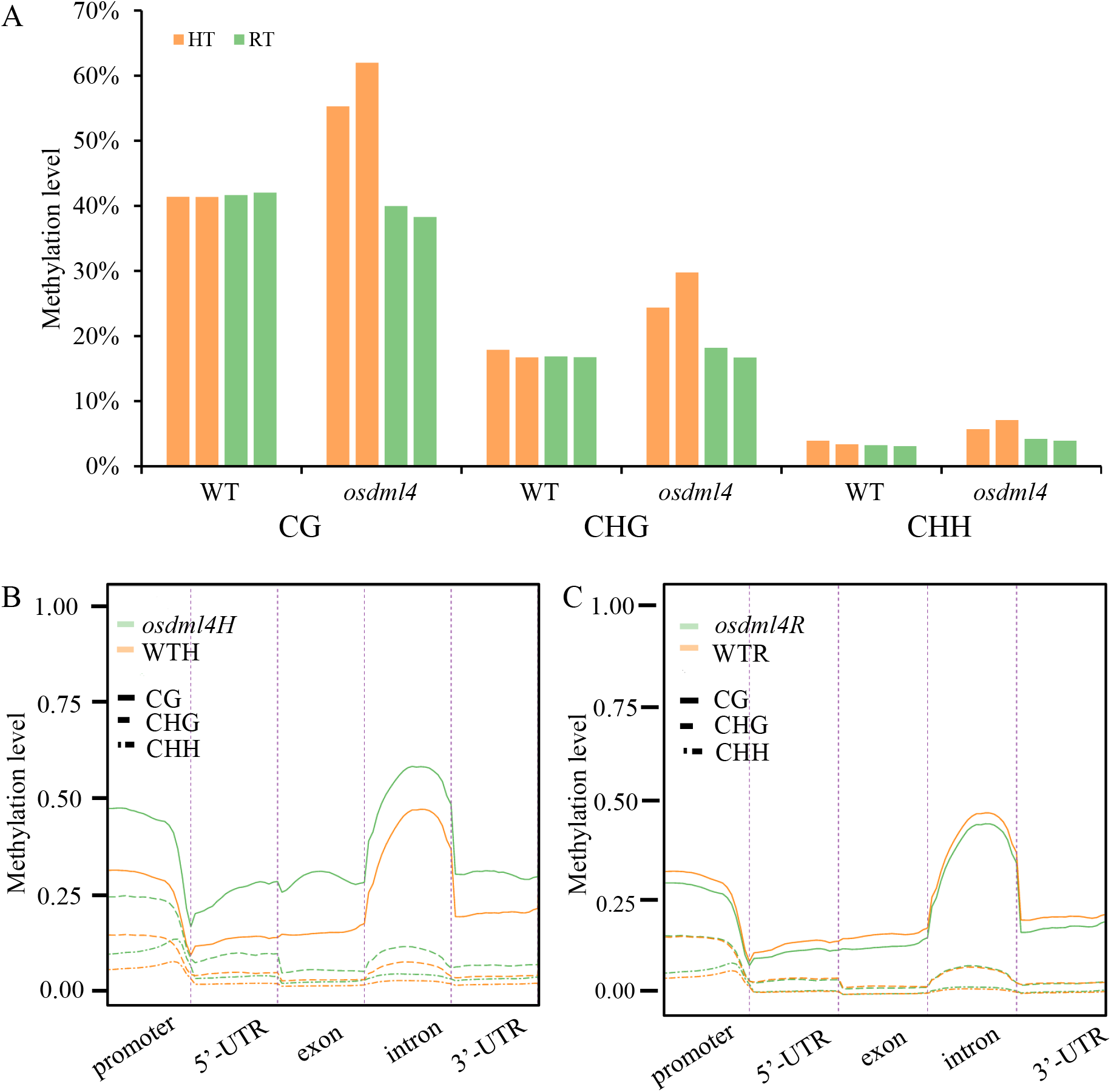
DNA methylation patter in 15 DAP seeds of WT and the *OsDML4* mutant under HT and RT. (A) DNA methylation levels in CG, CHG and CHH contexts in WT and the *osdml4* mutant seeds at 15 DAP under HT and RT. The average methylation levels in CG, CHG and CHH contexts of different genic regions in WT and the *osdml4* mutant seeds at 15 DAP under (B) HT and (C) RT.

The Integrated Genome Browser screenshots of the WGBS data showed that the DNA methylation levels of *RISBZ1* promoter (182 bp from −68 to −250 relative to the transcription start site) were dramatically hypermethylated under HT, while almost unchanged under RT. However, no obvious difference of the DNA methylation level was observed in the promoter of *RPBF*, glutelin genes, *VPE1*, *BiP1* and *PDIL1-1* under both HT and RT (Figure 8A-D, S4). The DNA methylation levels of *RISBZ1* and *RPBF* promoters were confirmed by Bisulfite Genomic Sequence (BSP) (Figure S5). These results suggested that the hypermethylation of *RISBZ1* promoter could directly reduce its expression, while others reduced gene expression was not directly affected by *OsDML4*-mediated DNA methylation.

**Figure 8.**
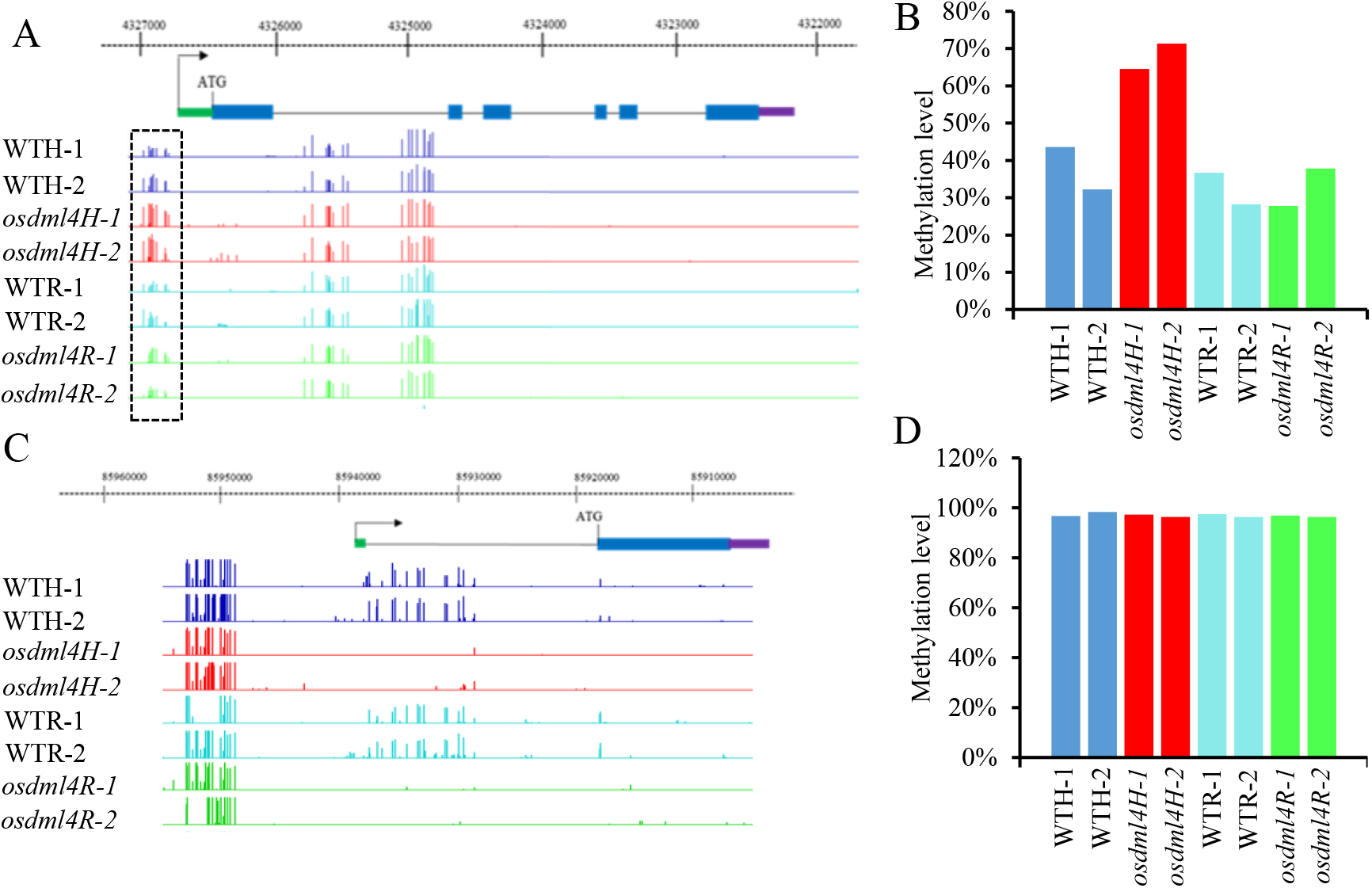
The DNA methylation levels in the *RISBZ1* and *RPBF*. The integrated genome browser screenshots of the WGBS data of the DNA methylation levels of (A) *RISBZ1* and (C) *RPBF* promoters, the statistic analysis of the DNA methylation levels of (C) *RISBZ1* and (D) *RPBF* promoters in WT and the *osdml4* mutant seeds at 15 DAP under HT and RT. Dashed frames indicate the hypermethylated regions in the *osdml4* mutants.

## Disscussion

### Knockout of *OsDML4* can induce floury endosperm

With global warming, the negative effects of HT on the growth and development of rice is getting increasingly obvious especially on the rice yield and quality (2-3). HT can increase the formation of chalky grains with a floury endpsperm (Nakata et al., 2017). Knockout mutants of *OsDML4* have a severe influence on the phenotype of the seeds with the floury endosperm only under HT (Figure 2A, B). Furthermore, the *osdml4* mutants had also effects on the seed storage materials, resulting in a decreased content of total starch and amylose, and an increase of total protein content (Figure 2D-F). The starch granules of the *osdml4* floury endosperm were irregularly round and loosely arranged, which was consistent with previous studies (Tabassum et al., 2020; Xu et al., 2020). These results suggested that knockout of *OsDML4* not only had an adverse effect on grain quality but also reduced grain yield under HT, indicating that the *OsDML4* gene plays a crucial role on the grain filling and is hypersensitive to HT.

### *OsDML4* can influence the accumulation of glutelins and the formation of PBs

The acccumulation of SSPs is closely related to formation of floury endosperm. In maize, a defective signal peptide in 22-kDa α-zein and the accumulation of the 24-kDa α-zein protein cause the floury endosperm phenotype (Coleman et al., 1997; Gillikin et al., 1997), αRNAi, combined βRNAi and γRNAi and the mutation of 16-kDa γ-zein also lead to opaque endosperm phenotypes (Kim et al., 2006; Wu and Messing, 2010). In rice, the grains of RNAi mutants with simultaneous supression of *GluA*, 13-kDa prolamin and globulin were opaque with a floury feature, resulted from less accumulation of glutelin A, 13-kDa prolamin, and globulin proteins and loosely packaged starch granules (Cho et al., 2016). HT will repress the deposition of the total starch and amylose to reduce the grain weight and yield during grain filling stage (Yamakawa and Hakata, 2010; Geigenberger, 2011; Sreenivasulu et al., 2015; Zhang et al., 2017). While the effect on SSPs, HT could reduce the acccumulation and expression of 13-kDa prolamin (Yamakawa et al., 2007), and increase the accumulation and the expression of glutelins (Lin et al., 2010; Cao et al., 2017). The loss-of-function *osdml4* mutants dramatically reduced the content of glutelins and 16-kDa prolamins, slightly reduced the 10-kDa prolamins, but no significantly affected the content of 13-kDa prolamins under HT (Figure 4A-B). The expression levels of SSP genes were consistent with the acccumulation of SSPs (Figure 5A), indicating that *OsDML4* plays an important role in SSPs transcription and protein acccumulation.

The formation of floury endosperm is usually accompanied by the disruption of the normal accumulation process of PBs (Wang et al., 2016; Chou et al., 2019; Ren et al., 2020). Knockout of *OsDML4* can significantly decrease the accumulation of glutelins and greatly reduce the number and the morphology of PB-IIs under HT. The incompleted PB-IIs with empty space were also found in the *osdml4* mutants, suggesting the existence of the restrain of vacuole formation (Figure 6A-F), which was consistent with the previous study (Kawakatsu et al., 2010). Moreover, *VPE1*, and *PDIL1-1* were significantly down-regulated in the *osdml4* mutant, indicating the *osdml4* mutant might suffer from ER stress to some extent (Figure 5B). The alteration of PBs is likely to be the key factor in the formation of floury endosperm.

### *OsDML4* is a new demethylase gene and whose function on cytosine demethylation depends on the temperature

DNA methylation represents one of the most important epigenetic regulotary mechanisms and plays a vital role in plant growth, development and response to biotic or abiotic sitmuli (Zhang et al., 2018). DNA methylation levels can be affected by abiotic stress conditions especially the temperature. Global disrupted DNA methylation induced by HT has a significant effect on microscope abortion and anther indehiscence in cotton (Ma et al., 2018). Active DNA demethylation is mainly controled by a series of transglucosylase gene family, including *DME*, *ROS1*, *DML2*, and *DML3*, the mutations of these genes can result in genome-wide hypermethylation (Gong et al., 2002; Hsieh et al., 2009; Qian et al., 2012; Yuan et al., 2020). We indentified a new demethylase gene *OsDML4* based on the conservative glycosylase domain. The WGBS analysis of 15 DAP seeds revealed that knockout of *OsDML4* can dramatically hypermethylate CG, CHG and CHH contexts in the whole genome under HT, while only a slightly increased methylation levels of CHG and CHH under RT (Figure 7), indicating that the demethylation function by *OsDML4* depends on the temperature. *RISBZ1* and *RPBF* are two essential TFs regulating SSPs exppression. Knock-down of *RPBF* or *RISBZ1* can only cause a sligthtly reduction of SSPs, whereas their double mutants resulted in a significant reduction of SSPs (Kawakatsu et al., 2009). Nevertheless, loss function of *OsbZIP58* significantly decreased SSPs under HT (Xu et al., 2020). In our study, we found that knockout of *OsDML4* can significantly repress the expression of *RPBF* and *RISBZ1* under HT, but not under RT (Figure5B, S3B). Furthemore, the methlation status of *RISBZ1* promoter was hypermethylated only under HT, while the methylation level of *RPBF* promoter was not changed (Figure 8), suggesting that *OsDML4* could regulate the expression of *RISBZ1* and *RPBF* to make opaque endersperm, but can only hypermethylate the *RISBZ1* promoter, not *RPBF* promoter. The mutation of *OsROS1a* to generate an extra transcript *mOsROS1a* with seven amino acids insertion exhibited thickened aleurones and the opague endosperm, which could result from DNA hypermethylation in the promoter regions of *RPBF* and *RISBZ1* to repress their expresson (Liu et al., 2018). Therefore, *OsDML4* could be a new demethylase gene that has different fuctions in demethylation from *OsROS1a* and the demethylation of *OsDML4* depends on the temperature.

In conclusion, knockout of *OsDML4* can genome-wide hypermethylate CG, CHG and CHH contexts under HT. The expression of two TFs *RISBZ1* and *RPBF* was reduced, but only accumulation levels of glutelins and that were not directly affected by DNA methylation. The reduced expression of *RISBZ1* under HT could be the hyper-methylation in its promoter, but the *RPBF* appears not to be affected directly by *OsDML4*-mediated DNA methylation.

## Materials and methods

### Plant materials and growth conditions

The rice cultivar Nipponbare was used to grow under natural field conditions with the daily mean day/night temperature 35 °C/26 °C as HT conditions during the booting stage. Before heading, the plants were transplanted to growth chamber with the day/night temperature 28 °C/22 °C as room temperature (RT) conditions. The immature seeds of 15 DAP (Days after pollination) were sampled for gene expression, PBs observation and whole-genome bisulfite sequencing (WGBS) analysis from both HT and RT conditions.

### Vector construction and rice transformation

The CRISPR/Cas9 binary vector pYLCRISPR/Cas9Pubi-H provided by Prof. Yaoguang Liu (South China Agriculture University) was used to construct target vectors for *OsDML4*. The online website (http://skl.scau.edu.cn/) was used to design two targets for *OsDML4* located at the first and the second exon of *OsDML4*, which were ligated to OsU6a and OsU6b promoters with the oligos of *OsDML4*-F1:5’-GCCGCAGTTCTCCGACTACGAGAC-3’ and *OsDML4*-R1:5’-AAAC GGTGGGGAACGCGGCCTTGA-3’ for target 1, and *OsDML4*-F2: 5’-GTTGACAGATCCCGCAAATACTCG-3’ and *OsDML4*-R2: 5’-AAACCGAGTATTTGCGGGATCTGT-3’ for target 2, respectively. The completed construct was introduced into *Agrobacterium tumefaciens* strain EHA105 by electroporation. Transgenic seedlings were obtained from regenerated hygromycin-resistant callus using selection medium containing 50 mg/L hygromycin and 500 mg/L cefotaxime. Genomic DNA was extracted from transgenic plants by CTAB method to detect the mutations.

### Scanning electron microscopy of starch granules

Mature rice grains were dried in an oven at 37 °C for 7 d and cooled in a drying apparatus. Cross-sections of the samples were manually fractured and sputter-coated with gold palladium on the surface. Magnifications of about 1000 and 3000 were used to observe endosperm and starch granule morphology with scanning electron microscopy (SEM) (JSM-6390LV).

### Transmission electron microscopic observation of protein bodies

Transverse sections (less than 1mm thick) of endosperms collected at the 15 DAP were fixed in 2.5% glutaraldehyde solution with 0.2 M phosphate buffer (pH7.2) for over 24 h. The sections were treated as described previously (Takemoto et al., 2002), embedded in Spurr’s low-viscosity and sectioned into ultra-thin sections. The ultra-thin sections were observed by transmission electron microscope (TEM) (Hitachi H7650).

### RNA Extraction and qRT-PCR Analysis

Spikelets were marked on the day of flowering and picked seed samples at 15 DAP. All seed samples were immediately put into liquid nitrogen for quick freezing and stored in −80 °C until use. Total RNAs of brown seeds were extracted by using the Plant DNA Mini Kit (Omaga) following the manufacturer’s protocol. The 1 μg total RNAs were used to synthesize cDNA with the PrimeScript™RT reagent Kit with gDNA Eraser (Takara). qRT-PCR was carried out with SYBR Premix Ex Taq II Premix (Takara) on the real-time system (Roche, Germany). Three replicates were set for each reaction, and β-*Actin* gene was used as the internal reference. The relative expressions were calculated by 2^−ΔΔCt^ method. All primers used are listed in Table S1.

### Determination of total protein, total starch and amylose content

The Kjeldahl method was used to determinate total protein content in rice, with slight modification according to the previous method (Kang et al., 2005). The 1 g of rice mature grain flour was put into a nitrate tube, and added 3 g of potassium sulfate: copper sulfate powder (10:1 w/w), then added 8 mL of concentrated sulfuric acid. The total nitrogen content was calculated based on the amount of hydrochloric acid consumed by the sample. Total protein content = total nitrogen content × 6.25. The total starch and amylose content were measured using the amylose and total starch assay kit (G0548W, Suzhou Grace Biotechnology Co., Ltd.)

### Extraction of glutelin and prolamin and SDS-PAGE

The extraction of each protein component was carried out according to the previous method (Takemoto et al., 2002) using 100 mg mature rice grain flour. Among them, the deionized water was used for albumin extraction, 2% NaCl (W/V) for globulin extraction, 70% ethanol for prolamin extraction, and 1% lactic acid for glutelin extraction. Glutelin and prolamin were separated by 12% SDS-PAGE.

### WGBS and analyses

Total genomic DNA was extracted from the immature 15 DAP seeds of the *osdml4* mutant and WT with two biological replicates using the CTAB method from both HT and RT conditions. A total amount of 100 ng genomic DNA spiked with 0.5 ng lambda DNA were fragmented by sonication to a mean size of 250 bp with Covaris S220. The DNA fragments were then treated with bisulfite using EZ DNA Methylation-GoldTM Kit (Zymo Research), and libraries were constructed by Novogene Corporation (Beijing, China). Subsequently, pair-end sequencing was performed using the Illumina Novaseq platform (Illumina, CA, USA).

### DNA methylation validated by Bisulfite Sequencing

Bisulfite sequencing PCR was performed using the same genomic DNA as for WGBS. In brief, 200 ng of DNA was treated with sodium bisulfite using the Qiagen Kit. The primers were designed using the MethPrimer online software (http://www.urogene.org/methprimer2/index.html) (Table S1). For each PCR reaction, 1 ul of bisulfite-treated DNA was used in a 20 ul reaction. PCR products were purified using Zymoclean Gel DNA Recovery kit and subcloned into pTA2 vector. For each DMR locus in the WT or the *osdml4* mutant, more than 10 independent clones were sequenced by Sanger method. The sequencing data were analyzed using the Bisulfite Analysis online software (http://katahdin.mssm.edu/kismeth/revpage.pl).

## Acknowledgements

This study was supported by the Talent Project of Zhejiang Province (2019R52033 to HMZ) and Zhejiang Zhengjingyuan Pharmacy Chain Co., Ltd. (H20151699 and H20151788 to JHX).

## Author contributions

JHX, HMZ and YY designed research; YY, CL, JRK, ZKY, JY, CCR, HMZ and JHX performed research; YY, CL, MSA conducted the field and growth chamber work; YY, ZL, JJZ, MHD, HMZ and JHX analyzed data; YY and JHX wrote and edited the manuscript.

## Supplemental Figure legends

**Figure S1. Phenotypes comparison of the *osdml4* mutants and WT at the mature stage under HT.** (A) Plant appearance of the *osdml4* mutants and WT. (B) Plant height of the *osdml4* mutants and WT. (C) Tiller numbers per plant of the *osdml4* mutants and WT. (D) Seed setting rate of the *osdml4* mutants and WT. (E) Seed numbers per panicle of the *osdml4* mutants and WT. Data are the means ± SD of three biological replicates. Significant differences were determined using two-tailed Student’s t-test (*p<0.05, **p<0.01).

**Figure S2. Appearance of the WT and *osdml4* mutants mature seeds under RT.** (A) External appearance of brown seeds. (B) Transverse sections of the dry seeds.

**Figure S3. Expression levels of genes related to seed storage proteins biosynthesis (A) and transmission electron microscope (TEM) of protein bodies in developing endosperm at 15 DAF under RT (B-E).** B and C, WT; D and E, *osdml4* mutants. Scale bars=10 μm in (A, B) and 2μm in (C, D).

**Figure S4. The DNA methylation levels in the promoter regions of glutelin genes (*GluA1, GluB1, GluC*, and *GluD*) and their regulatory factors (*VPE1, BiP1* and *PDIL1-1*) in WT and *osdml4* mutants under HT and RT at 15 DAP.**

**Figure S5. DNA methylation levels of *RISBZ1* and *RPBF* promoters obtained by BSP.**

**Supplemental Table 1 The sequences of primers used in this study**

**Supplemental Table 2 Details of data quality control statistics of WGBS sequencing generated libraries**

